# Sec23b regulates cell migration by orchestrating collagen I secretion and processing

**DOI:** 10.64898/2026.03.04.709595

**Authors:** E. Emily Joo, Audrey Astori, Jonathan St-Germain, Brian Raught, Michael F. Olson

## Abstract

Cell migration is critically important for development and for homeostatic processes such as wound healing, while its aberrant regulation may contribute to the development of pathological conditions and diseases. Efficient cell migration requires the coordination of many structural and signalling proteins (*e.g.* Rho GTPases) that have key functions in controlling actin cytoskeleton organization and dynamics. Given that the actin cytoskeleton has a central role in orchestrating the complex series of events underlying cell migration, characterizing the actin-associated proteome can illuminate important molecular players that could be novel therapeutic targets for diseases associated with dysregulated cell migration. Here, we report a proteomic-based assay that uses a doxycycline-inducible proximity ligation assay to label actin-associated proteins (actin cytoskeletome) during the early stages of cell migration. We found that an unexpected actin-associated protein, Sec23b, influences the rate of cell migration by orchestrating the secretion and post-secretion maturation of Collagen I.

## Introduction

Cell migration is a fundamental biological process necessary for the normal growth and health of all living organisms^1–4^. Aberrant cell migration may lead to developmental disorders and perturbations in homeostatic mechanisms that can contribute to pathological conditions including chronic inflammation, atherosclerosis and cancer metastasis^5–8^. Cancer metastasis is a multi-step process that starts with cells dissociating from primary tumours, which then migrate to distant locations via blood and lymphatic vessels, and ultimately colonize at secondary sites^9–12^. Although the precise mechanisms and molecular players that contribute to cancer metastasis have yet to be fully characterized, it is clear that altered cell migration is a hallmarks of solid tumour metastasis^9–11,13^.

Many different cell types migrate through various heterogeneous environments in response to migratory cues as part of processes including normal development, wound healing and inflammatory responses^1,7,14,15^. Numerous signalling pathways and protein complexes play crucial roles in cell migration. Whether the migratory behaviour is guided or stochastic, collective or individual, the migration of mesenchymal or mesenchymal-like cells on 2-dimensional surfaces can be categorized into several stages. Initially, cells extend specialized protrusions such as ruffles, lamellipodia and filopodia to move forward and actively survey their immediate microenvironment, an activity primarily dependent on dynamic reorganization of actin filaments^1,16,17^. For efficient migration, these exploratory outgrowths typically establish new integrin-dependent or independent contacts for cells to attach onto the extracellular matrix^18–21^. These newly formed cell-matrix anchor points are key constituents of the leading edge, establishing the subsequent forward displacement of cellular components, such as the nucleus and organelles, during cell migration. Finally, cells detach the trailing end by forcefully pulling the long-lived cell-matrix adhesions and/or internalizing these attachments. These coordinated activities propel the forward motion of the cell^1–4^.

Of the numerous molecular players in cell migration and cancer metastasis, the actin cytoskeleton is arguably the foremost^2,18^. The actin cytoskeleton is one of the major cytoskeletal systems in cells, with actin constantly cycling between filamentous (F-actin) and globular (G-actin) states. Given the importance of regulating the balance between these two states, many previous research efforts have sought to identify actin-binding proteins that regulate actin dynamics and organization in various cellular and subcellular locations. For example, protein-protein interaction assays, including immunoprecipitation, yeast 2-hybrid, and GST-protein pulldowns, have been used to identify actin-binding proteins^19–25^. Additionally, chemical cross-linkers have been used to identify and characterize actin-interacting proteins^26,27^. However, these methods primarily characterize protein-protein interactions at steady state. Although microscopy-based assays have revealed information on the spatial and temporal regulation of actin dynamics by the Rho family GTPases^28–30^, these studies are only able to examine candidate proteins on a case-by-case basis to characterize their roles in regulating the actin cytoskeleton^28–30^.

To acquire an expanded view of the dynamic actin cytoskeletome, we combined the 17 amino acid Lifeact peptide, derived from the N-terminus of the *Saccharomyces cerevisiae* actin binding protein ABP140^31^, with the improved BirA biotin ligase (miniTurboID)^32^ for highly sensitive and rapid labelling of nearby proteins. The miniTurboID-Lifeact fusion protein was placed under the transcriptional control of a doxycycline-inducible promoter. Here, we report that miniTurboID-Lifeact identified differentially labelled proteins in response to scratch-induced migration, including the F-actin binding proteins Moesin and α-Actinin 4 as well as the focal adhesion proteins Vinculin and LIM-containing lipoma-preferred partner (LPP). In addition, Sec23b, a component of the COPII coat complex^33^, was unexpectedly identified as being more labelled by miniTurboID-Lifeact following scratch wounding. Knockdown experiments demonstrated that Sec23b promotes efficient migration by governing the secretion and proteolytic processing of secreted collagen I. Together, these findings indicate a previously unknown role for Sec23b in regulating extracellular matrix organization to enable efficient cell migration.

## Results

### MiniTurboID-Lifeact probe labelling actin cytoskeletome during scratch-induced migration

We previously created a FLAG-epitope tagged BioID-Lifeact proximity ligation probe that could be used to identify changes in actin binding proteins during various cellular processes^34^. This probe was improved by replacing the N-terminal BirA with the smaller and more active miniTurbo^32^ version, and its transcription was placed under the control of a doxycycline-inducible promotor. The expression of miniTurboID-Lifeact in the MDA-MB-231 triple-negative metastatic breast cancer cell line was induced with the addition of 25 nM doxycycline for 18 hours. The expressed miniTurboID-Lifeact localized with endogenous actin structures and mediated protein biotinylation that was detected with fluorescently-labelled streptavidin, but not observed in control cells (**Figure 1A**). Moreover, miniTurboID-Lifeact expression did not affect the closure of scratches made in confluent MDA-MB-231 monolayers relative to control cells (**Figure S1A**), indicating that there were no detectable deleterious effects on cell migration. Given that Lifeact has higher affinity for G-actin relative to F-actin^31^, we sought to determine if miniTurboID-Lifeact could detect changes in proteins associated with F-actin structures when they were disrupted by Cytochalasin D. After doxycycline induction, miniTurboID-Lifeact expressing MDA-MB-231 cells were treated for 1 h with 5 µM Cytochalsin D or an equal volume of vehicle control (DMSO, control), followed by a 50 µM biotin pulse for 30 minutes to label proximal proteins with biotin. Biotinylated protein enrichment with streptavidin-conjugated beads followed by mass spectrometry identified a total of 171 proteins that had Bayesian false discovery rates (BFDR) < 0.05 in Cytochalasin D treated cells, of which 36 proteins had ≥ 2 fold reduced biotin labelling and 34 proteins had ≥ 2 fold increased biotin labelling (**Figure 1B. Table S1**). Gene Set Enrichment Analysis (GSEA)^35^ of the proteins’ molecular functions only identified 2 in common for both sets of reduced and increased biotinylated proteins; “actin binding” and “cytoskeleton protein binding” (**Figure 1C, Tables S2-S3**). The third highest ranked molecular function for the reduced biotinylated protein set was “actin filament binding”, which was not identified with the increased biotinylated protein set. These results are consistent with the increased biotin labelled proteins being predominantly G-actin binding proteins, and reduced biotin labelled proteins being predominantly F-actin binding proteins, concordant with the effect of Cytochalasin D being to block actin polymerization and shift the cellular actin pool away from F-actin and toward G-actin. STRING protein-protein association database^36^ mapping of the 27 proteins (19 reduced biotinylation + 8 increased biotinylation) grouped in the “actin binding” molecular function plus β-actin revealed a network of interactions between increased biotinylated (green symbols) and decreased biotinylated (red symbols) proteins with β-actin (blue symbol) at the centre (**Figure 1D**). Well characterized G-actin binding proteins in the increased biotinylated group that were linked with actin in the STRING network^36^, include Profilin1 (PFN1)^37^, Diaphanous1 (DIAPH1)^38^, ENA (ENAH)^39^, JMY^40^ and Spire1^41^. F-actin binding proteins in the decreased biotinylated group linked with actin in the STRING network include the F-actin binding proteins Drebin1 (DBN1)^42^, Supervillin (SVIL)^43^, Synaptopodin (SYNPO)^44^, Caldesmon 1 (CALD1)^45^ and Nexilin (NEXN)^46^, as well as the focal adhesion proteins Vinculin (VCL) and Moesin (MSN)^47^. Taken together, these observations validate miniTurboID-Lifeact as a useful tool to identify dynamic changes in actin-associated protein complexes.

**Figure 1.**
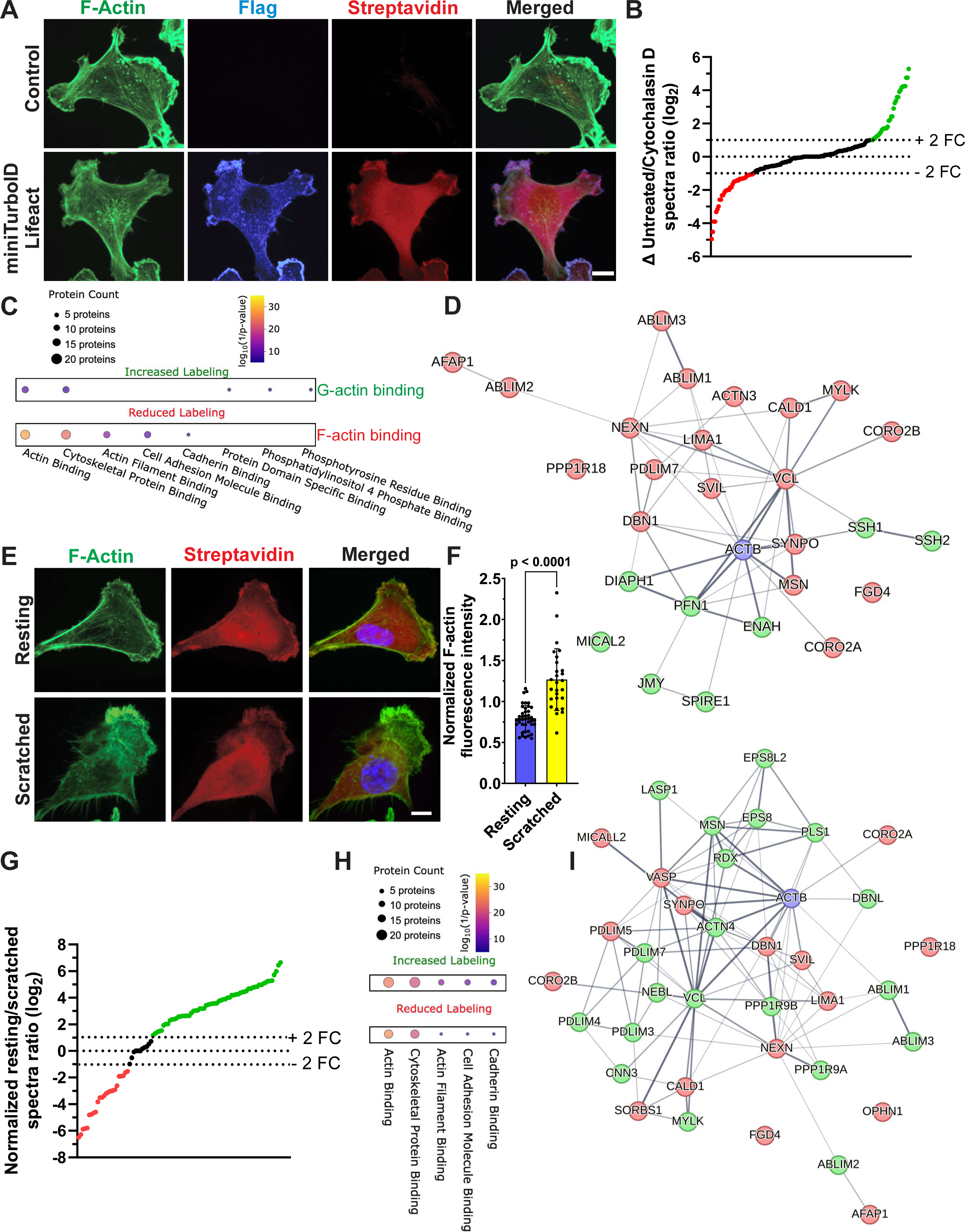
MiniTurboID-Lifeact identification of dynamic changes in the actin interactome. **A.** Immunofluorescence images of control or miniTurboID-Lifeact expressing MDA-MB-231 human breast cancer cells stained for F-actin (green), FLAG-tagged miniTurboID-Lifeact (blue) or streptavidin labelled protein biotinylation (red). Scale bar = 10 µm. **B.** Log_2_ change in protein spectra ratio between untreated and Cytochalasin D treated miniTurboID-Lifeact expressing MDA-MB-231 cells. Dotted lines indicate the division between proteins that were less biotinylated (red) by two or more fold change (-2 FC) or more biotinylated (green) by two or more fold change (+ 2 FC). **C.** Gene set enrichment analysis (GSEA) of the top five most statistically significant molecular functions of the 36 proteins with Bayesian false discovery rates (BFDR) < 0.05 and ≥ 2 fold reduced biotin labelling, and 34 proteins with FDR < 0.05 and ≥ 2 fold increased biotin labelling in Cytochalasin D treated cells. **D.** STRING protein-protein network mapping of the 27 proteins (19 reduced biotinylation (red) + 8 increased biotinylation (green)) grouped in the GSEA “actin binding” molecular function plus β-actin (blue). **E.** MDA-MB-231 cells expressing miniTurboID-Lifeact were either unscratched (resting) or scratched to remove ∼2/3 of the cells to initiate cell migration. After 1.5 h, cells were fixed and stained for F-actin (green) and streptavidin labelled protein biotinylation (red). Scale bar = 10 µm. **F.** Mean normalized phalloidin fluorescence intensity for resting (n = 41) and scratched (n = 27) cells. Unpaired student’s t-test. Means ± standard deviation. **G.** Log_2_ resting/scratched protein spectra ratio of proteins with a BFDR < 0.05 in the scratched condition. Dotted lines indicate the division between proteins that were less biotinylated (red) by two or more fold change (-2 FC) or more biotinylated (green) by two or more fold change (+ 2 FC). **H.** GSEA of the top five most statistically significant molecular functions of the 61 proteins that had BFDR < 0.05 and ≥ 2 fold increased biotinylation, and 24 proteins that had BFDR < 0.05 and ≥ 2 fold decreased biotinylation. **I.** STRING protein-protein network mapping of the 36 proteins (20 reduced biotinylation (red) + 16 increased biotinylation (green)) grouped in the GSEA “actin binding” molecular function plus β-actin (blue).

To identify changes in actin-associated protein complexes during the early initiation phase of cell movement, uniformly distributed scratch wounds that removed ∼⅔ of cells from doxycycline-induced confluent MDA-MB-231 monolayers were made to induce actin cytoskeleton reorganization associated with widespread migration into the scratched areas. Cells were either unscratched (resting) or scratched, left for 1 hour and then pulsed with 50 µM biotin for 30 minutes to label actin-associated protein complexes (**Figure 1E**). Quantification of mean phalloidin fluorescence intensity in resting or scratched cells showed significantly > 50% increased F-actin levels in migrating cells (**Figure 1F**). Time-lapse analysis revealed that 1.5 hours after scratch wounding was sufficient for protrusions to form and for cells to start migrating into scratched areas, but before they migrated a distance greater than a full cell length (**Figure S1B**). Biotinylated proteins from resting and scratched conditions were enriched with streptavidin beads and subjected to mass spectrometry for protein identification. The ratios of peptide spectra for each identified protein relative to auto-biotinylated miniTurboID-Lifeact were compared in each condition. Fold changes were calculated by normalizing the difference in the ratio of each protein’s peptide spectra to the miniTurboID-Lifeact spectra between resting and scratched cells relative to the mean difference for all proteins. Using a BFDR < 0.05 cut-off in the scratched cell condition to filter proteins based on the confidence of their identification, 61 proteins had ≥ 2 fold increased biotinylation and 24 proteins had ≥ 2 fold decreased biotinylation (**Figure 1G, Table S4**). GSEA identified the same top 5 molecular functions in the same rank order for the increased and decreased biotinylated protein subsets (**Figure 1H, Tables S5-S6**), with “actin binding” being the highest ranked. STRING network mapping of the 36 proteins (20 increased biotinylation + 16 decreased biotinylation) grouped in the “actin binding” molecular function plus β-actin, which was added to facilitate the visualization, revealed a network of interactions between increased biotinylated (green symbols) and decreased biotinylated (red symbols) proteins with β-actin (blue symbol) at the centre (**Figure 1I**). One prediction is that proteins that were less biotinylated in Cytochalasin D treated cells in which F-actin was disrupted (**Figure 1B, Table S1**) would be more biotinylated in scratched cells (**Figure 1G, Table S4**) in which F-actin levels were increased (**Figure 1F**). Comparison of the “actin binding” molecular function STRING protein networks in **Figures 1D and 1I** reveal some proteins in agreement with this prediction, including Vinculin (VCL), Moesin (MSN)^47^, Acting binding LIM protein 1 (ABLIM1)^48^, ABLIM2^49^, ABLIM3^50^, Myosin light chain kinase (MYLT)^51^ and PDZ and LIM protein 7 (PDLIM7, also known as Enigma)^52^. However, several actin-binding proteins had lower biotinylation in both Cytochalasin D treated cells (**Figure 1D**) and in scratched cells compared to controls (**Figure 1I**), including Actin filament-associated protein 1 (AFAP1)^53^, Coronin 2A (CORO2A)^54^, Coronin 2B (CORO2B)^55^, Drebin1 (DBN1)^56^, Supervillin (SVIL)^43^, Synaptopodin (SYNPO)^44^, Caldesmon 1 (CALD1)^45^, Nexilin (NEXN)^46^, LIM domain and actin-binding protein 1 (LIMA1, also known as EPLIN)^57^, Phostensin (PPP1R18)^58^ and FYVE, RhoGEF and PH domain containing 4 (FGD4, also known as Frabin)^59^, suggesting reduced association with F-actin as the cytoskeleton was re-organized during the initial stages of cell migration induced by scratching. There were no actin-binding proteins with increased biotinylation in Cytochalasin D (**Figure 1D**) treated cells that were either more or less biotinylated in scratched cells (**Figure 1I**), possibly indicating that the total cellular pool of G-actin was little changed during the early stages of scratch-induced cell motility. Taken together, these observations indicate that there are changes in the actin-associated proteome as cells transition from static to migrating that are both dependent and independent of the increased levels of F-actin, likely reflecting changes in protein-protein interactions that contribute to cytoskeleton reorganization in addition to changes in actin polymerization.

When the top 10 increased biotinylated proteins in scratched cells were evaluated by GSEA, 7 were categorized in the “actin binding” molecular function group: Drebin-like (DBNL), Calponin 3 (CNN3), α-actinin 4 (ACTN4), ABLIM3, PDLIM4, PDLIM7 and Moesin (MSN). The second most highly biotinylated protein LIM-containing lipoma-preferred partner (LPP) is placed in the Gene Ontology Biological Process (GOBP) gene set “cell adhesion” and is a scaffold that interacts with proteins including α-Actinin, LASP1, VASP and Supervillin to promote cell migration^60^. The seventh most biotinylated protein SH3 And PX Domains 2B (SH3PXD2B, also known as TKS4) is in the GOBP gene set “extracellular matrix disassembly” and also is a scaffold that interacts with Cortactin and Src to enable membrane ruffling, lamellipodia formation, cell motility^61^ and the formation of extracellular matrix degrading invadopodia structures ^62^. The outlier was the third most highly biotinylated protein Sec23b, which is a core component of the coat protein complex II (COPII) that functions to transport newly synthesized proteins that are destined to intra- or extracellular space from the endoplasmic reticulum (ER) to the Golgi apparatus^33,63^. Sec23b was also less biotinylated in Cytochalasin D treated cells (-1.9 FC) as determined by mass spectrometry, consistent with its proximity to F-actin structures.

### Sec23b associates with microtubules but switches to filamentous actin during the early phases of cell migration

To validate our proteomic findings, we assayed the changes in Sec23b biotinylation in two ways. Firstly, in addition to the FLAG-epitope tagged N-terminal miniTurboID-Lifeact fusion protein, a doxycycline-inducible FLAG-epitope tagged C-terminal Lifeact-miniTurboID fusion construct was generated and expressed in MDA-MB-231 cells (**Figure S1C**). Secondly, actin filaments in miniTurboID-Lifeact or Lifeact-miniTurboID expressing cells were disrupted by treatment with Cytochalasin D (CytoD) for 1 hour. Expression of the FLAG-epitope tagged miniTurboID-Lifeact or Lifeact-miniTurboID fusion proteins did not vary in each condition (**Figure 2A, top bands**). When biotinylated proteins were enriched with streptavidin-conjugated beads and then western blotted, there was comparable biotinylation of the intermediate filament protein Vimentin in all conditions (**Figure 2A**). In agreement with the mass spectrometry results that showed increased biotinylation of Moesin and Sec23b in scratch-wounded miniTurboID-Lifeact expressing cells, both Moesin and Sec23b were biotinylated to a greater extent in scratch-induced miniTurboID-Lifeact and Lifeact-miniTurboID expressing cells (**Figure 2A**). Furthermore, Moesin and Sec23b biotinylation were decreased in CytoD treated cells expressing either miniTurboID-Lifeact or Lifeact-miniTurboID (**Figure 2A**). Thus, these results support the conclusion that scratch-induced cell mobilization (**Figure S1B**) and accompanying cytoskeleton reorganization (**Figure 1F**) promoted the association of Moesin and Sec23b with the actin cytoskeleton.

**Figure 2.**
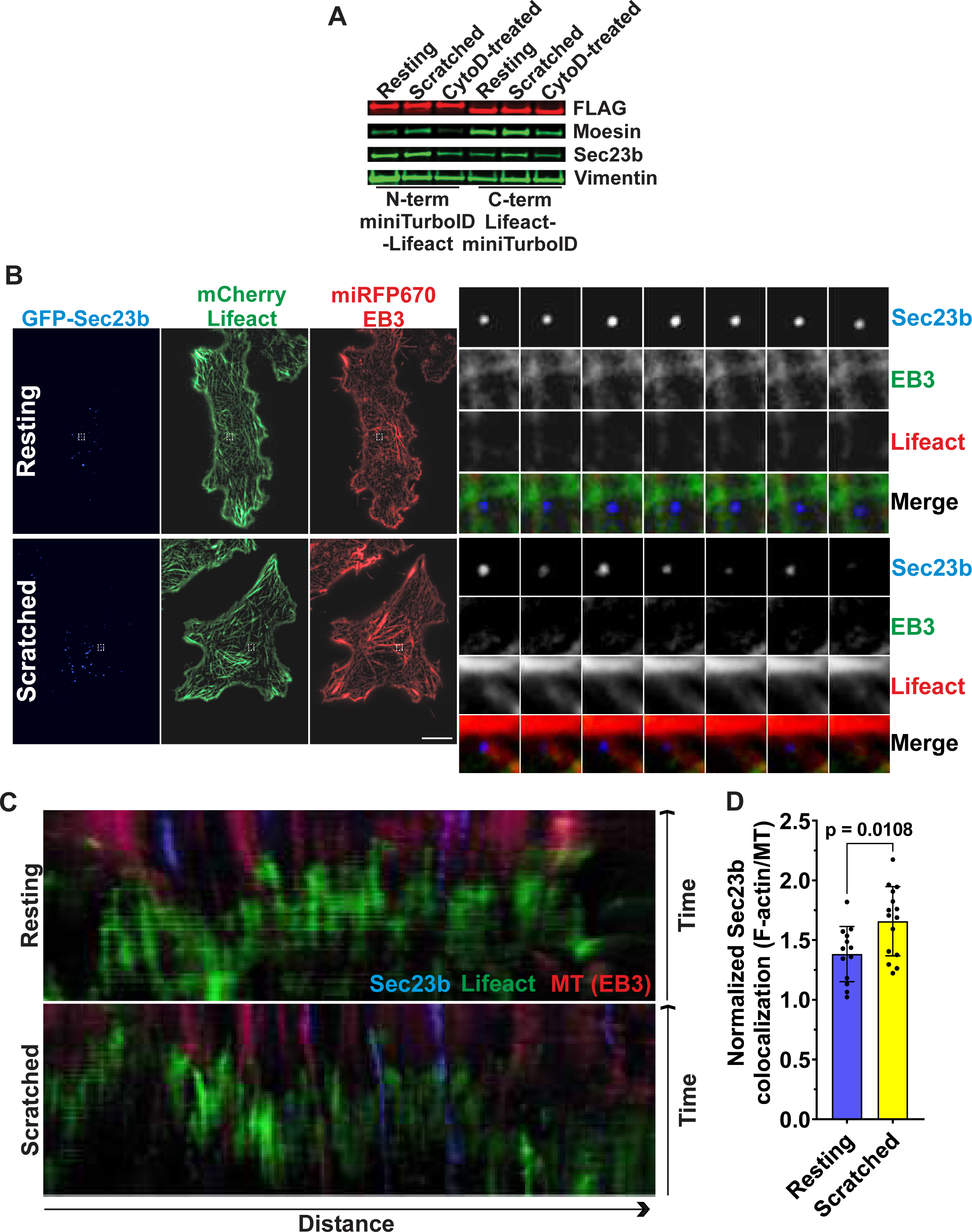
Sec23b localization during resting and early phases of migration. **A.** Western blot of streptavidin pulldown from MDA-MB-231 cells expressing miniTurboID-Lifeact under indicated conditions. **B.** Still images of MDA-MB-231 cells expressing GFP-Sec23b, mCherry-Lifeact or 670-EB3. Time montage of the boxed area is magnified and shown on the right side. **C.** Representative kymographs of time-lapse images of MDA-MB-231 cells expressing GFP-Sec23bB, mCherry-Lifeact or 670-EB3. For visualization, Sec23bB is pseudocoloured blue, mCherry-Lifeact is pseudocoloured green and 670-EB3 is pseudocoloured red. **D.** Normalized Sec23b colocalization ratio for F-actin relative to microtubules (MT) for resting (n = 13) and scratched cells (n = 15). Unpaired student’s t-test. Means ± standard deviation.

The observation of increased miniTurboID-Lifeact mediated Sec23b biotinylation in migrating cells was unexpected because Sec23b is a component of the COPII coat protein complex composed of Sar1, Sec23a/Sec23b, Sec24, Sec13 and Sec31, which is responsible for vesicle budding from the endoplasmic reticulum (ER) to transport proteins to the Golgi complex^63^. COPII vesicles can interact with the dynactin-dynein motor complex to enable microtubule-mediated vesicle transport^33^. Given that the microtubule network and actin cytoskeleton are highly dynamic^64–67^, we tracked Sec23b localization by live-cell imaging. To visualize Sec23b, microtubules and the actin cytoskeleton simultaneously in living cells, we tagged Sec23b with GFP (GFP-Sec23b), identified microtubules with EB3 fused with 670 fluorescent protein (670-EB3), and labelled the actin cytoskeleton with Lifeact tagged with mCherry fluorescent protein (mCherry-Lifeact). These fluorescently tagged proteins have been extensively used in numerous live imaging studies and their representations of the target proteins have been well validated^31,68–71^, and cells were selected by flow cytometry with expression below median level levels to minimize the risk of mislocalization or undesired effects. As shown in representative stills from live-cell time-lapse imaging (**Figure 2B**), Sec23b puncta typically appeared to localize in the vicinity of EB3-positive microtubules in resting unscratched cells. In contrast, Sec23b particles shifted towards Lifeact-labelled F-actin when cells were induced to migrate into scratched areas (**Figure 2B**). The shift from the EB3-positive microtubule network to Lifeact marked F-actin in migrating cells could be also observed by visualizing Sec23b particles, actin filaments and microtubule networks using kymographs. As shown in **Figures 2C**, Sec23b moved from the microtubule network to F-actin when MDA-MB-231 cells were induced to move into scratched areas. When the colocalization between Sec23b with actin versus microtubules was compared for the resting/equilibrium state versus the scratched/migratory state, there was a significant shift from microtubules to actin filaments (**Figure 2D**), indicating that a proportion of Sec23b switches from associating with microtubules to actin filaments during scratch-induced cell migration.

### Depletion of Sec23b inhibits MDA-MB-231 cell spreading and migration

The observation that scratch-induced cytoskeleton reorganization (**Figure 1F**) was paralleled by increased miniTurboID-Lifeact mediated Sec23b biotinylation (**Figure 2A**) suggested that it might play a role in the initial stages of cell migration. To examine the role of Sec23b in cell migration, it was knocked down by expressing two independent shRNAs. As shown in **Figure 3A**, KD1 or KD2 shRNAs depleted Sec23b by >50% in MDA-MB-231 cells compared to the scrambled shRNA control sequence. Sec23b knockdown in MDA-MB-231 cells by either shRNA did not induce significant differences in single cell F-actin fluorescence intensity relative to wheat germ agglutin (WGA) whole cell staining (**Figure 3B)** or F-actin organization as determined by measuring actin fibre anisotropy (**Figure 3C**) as visualized by indirect staining with fluorescent phalloidin. These results are consistent with Sec23b not having a direct role in regulating actin cytoskeleton organization. Given that the efficacy of mesenchymal cell migration on 2-dimensional surfaces is intimately related to cell spreading ^50^, we measured spread cell areas over time. As shown in **Figure 3D**, MDA-MB-231 cells with reduced levels of Sec23b protein were significantly ∼ 20% relatively less spread compared to control cells 2.5 hours after plating. To determine if there were any effects of Sec23b knockdown on how cells interacted with the underlying substrate, focal adhesions in control and Sec23b knockdown cells were visualized by staining with phalloidin and anti-Paxillin antibody (**Figure 3E**). There was a small but statistically significant decrease in the average elliptical form factor in the Sec23b knockdown cells (**Figure 3F**), indicating a trend of rounder focal adhesions. In addition, quantitative analysis of focal adhesion sizes^72–74^ revealed that knockdown cells had an increased proportion of smaller focal adhesions (0 to 1.5%, **Figure 3G**) and decreased portion of larger focal adhesions (1.5-3%, **Figure 3H**). Sec23b knockdown also significantly delayed scratch wound closure (**Figures 3I and 3J**) and significantly reduced random cell migration velocities (**Figures 3K and S2A-B**) without significantly altering turning angle vectors (**Figure S2C**) or migration directionality (**Figure S2D**). Taken together, these observations indicate that Sec23b contributes to efficient cell migration, at least partially mediated by enabling cell spreading and focal adhesion assembly/maturation.

**Figure 3.**
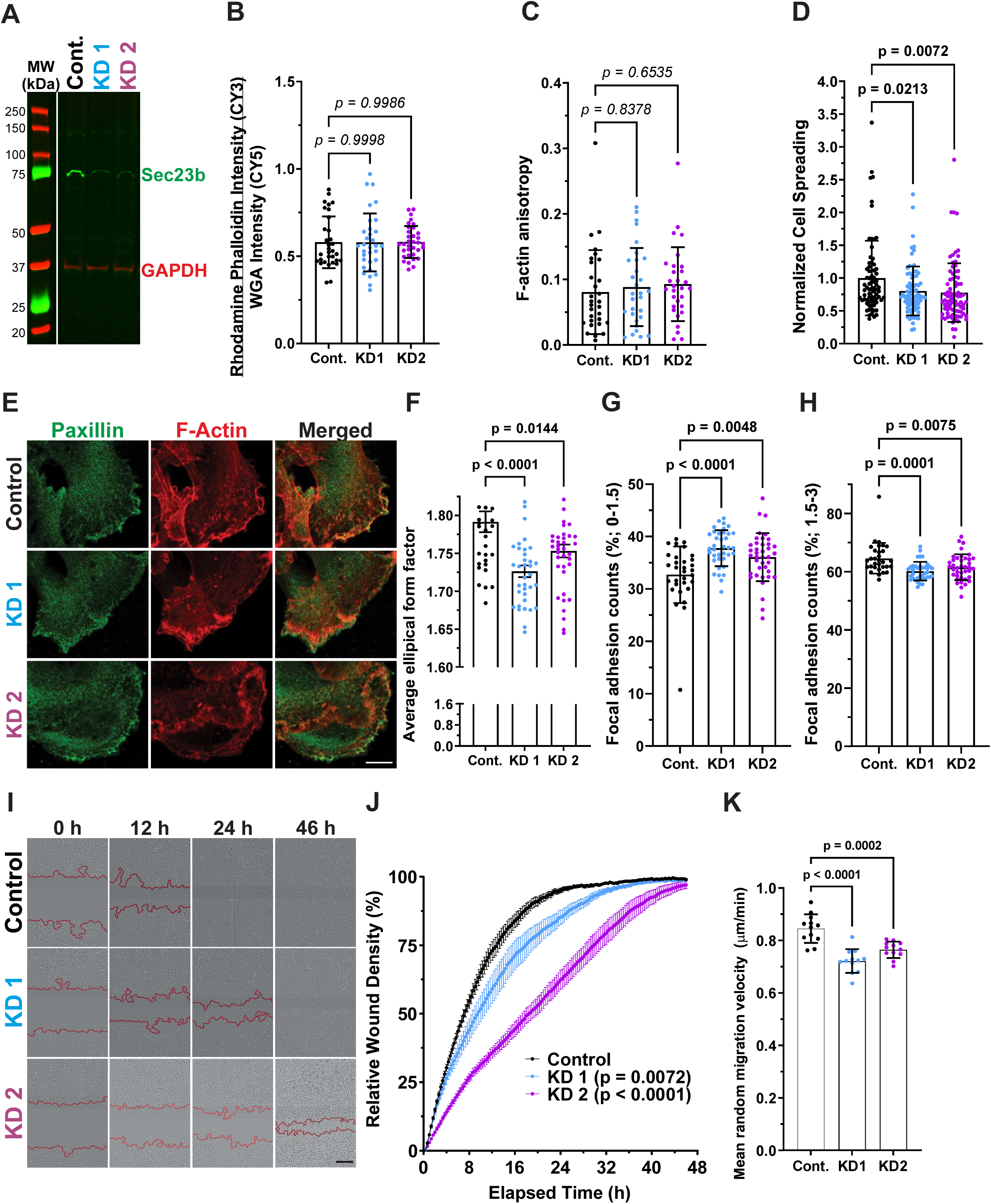
Sec23B depletion decreases scratch-induced and random migration. **A.** Western blot of MDA-MB-231 cells expressing control scrambled shRNA (Cont.) or Sec23b shRNAs (KD1 and KD2). Each cell line was induced with 2 µM doxycycline for 3 days. **B.** Relative fluorescence intensity of rhodamine-conjugated phalloidin relative to Cy5-conjugated wheat germ agglutinin (WGA) whole cell stain for Cont. (n = 32), KD1 (n = 32) and KD2 (n = 34) cells. One way ANOVA with Dunnett’s multiple comparison test. Means ± standard deviation. **C.** Relative F-actin anisotropy for Cont., KD1 and KD2 cells (n = 30). One way ANOVA with Dunnett’s multiple comparison test. Means ± standard deviation. **D.** Normalized cell spread areas for Cont. (n = 72), KD1 (n = 76) and KD2 (n = 82) cells. One way ANOVA with Dunnett’s multiple comparison test. Means ± standard deviation. **E.** Immunofluorescence images of MDA-MB-231 Cont., KD1 or KD2 cells that were fixed and stained with paxillin antibodies and rhodamine phalloidin. Scale bar = 10 μm. **F.** Elliptical factor form (a measurement of length over width) of each paxillin-positive adhesion for Cont. (n = 31), KD1 (n = 37) or KD2 (n = 39) cells. One way ANOVA with Dunnett’s multiple comparison test. Means ± standard deviation. **G,H.** Proportion of paxillin-positive adhesions in Cont. (n = 31), KD1 (n = 37) or KD2 (n = 39) cells arbitrary grouped based on their sizes as 0-1.5 (small size) and 1.5-3 (medium size). One way ANOVA with Dunnett’s multiple comparison test. Means ± standard deviation. **I.** Scratch-induced migration of MDA-MB-231 cells expressing control scrambled shRNA or Sec23b shRNAs KD1 or KD2. Scale bar = 300 μm. **J.** Relative wound densities for Control, KD1 and KD2 cells (n = 12) over 48 h. Nonparametric Kruskal-Wallis multiple comparison test. Means ± standard error. **K.** Mean random cell migration velocities of random cell migration for Cont. (n = 31), KD1 (n = 37) or KD2 (n = 39) cells. One way ANOVA with Dunnett’s multiple comparison test. Means ± standard deviation.

### Sec23b is important for collagen I secretion and processing, impacting cell migration

Given that Sec23b is a component of COPII mediated vesicle movement, we wondered whether the movement of COPII mediated vesicle movements would be altered when Sec23b was depleted. As shown in **Figure 4A**, we tracked the movement of COPII vesicles marked by glycosylphosphatidylinositol (GPI) anchors, a common marker used to assay COPII-mediated trafficking^75–77^, and found a significant ∼2-fold increase in the rate of trafficking in cells depleted of Sec23b, suggesting that COPII-mediated vesicle trafficking was aberrantly regulated. Further, we hypothesized that depletion of Sec23b would affect collagen I secretion because the COPII pathway has been reported to facilitate collagen I trafficking^33,78–83^ and cancer cells secrete extracellular matrix (ECM) proteins^84–86^. Newly synthesized collagen monomers heterotrimerize into large triple helices (procollagen I) that are packaged into COPII mediated vesicles to enable the bulky procollagen I to be transported from ER to Golgi apparatus, and finally to be secreted out from cells^87–89^. Once secreted into the extracellular space, procollagen I must be cleaved at the N- and C-termini (telocollagen I) to then be assembled into insoluble collagen filaments, a physiological component of extracellular matrices^90,91^. Thus, it was unexpected to observe a significant > 20-fold increase of the secreted soluble procollagen I in conditioned culture media in Sec23b knockdown cells, as detected with an antibody raised against the N-terminal procollagen domain (**Figures 4B and 4C**). To ensure correct handling and normalization of the conditioned media, recombinant His-tagged MRCKβ protein was added as a sample preparation marker to equivalent volumes of conditioned media from each condition (**Figures 4B and 4C**). Interestingly, the increase in secreted procollagen I did not correspond to a concomitant increase in proteolytically cleaved collagen monomers (telocollagen I), a marker of physiologically active collagen I filaments^87,92^. Instead, a significant ∼25% decrease in biologically active telocollagen I in Sec23b knockdown MDA-MB-231 cells was observed by quantifying telocollagen fibres using immunofluorescence microscopy (**Figures 4D and 4E**). Taken together, these observations indicate that Sec23b depletion resulted in increased secretion of procollagen I proteins into the extracellular space with a concomitant decreased level of physiologically active collagen I filaments deposited in the ECM.

**Figure 4.**
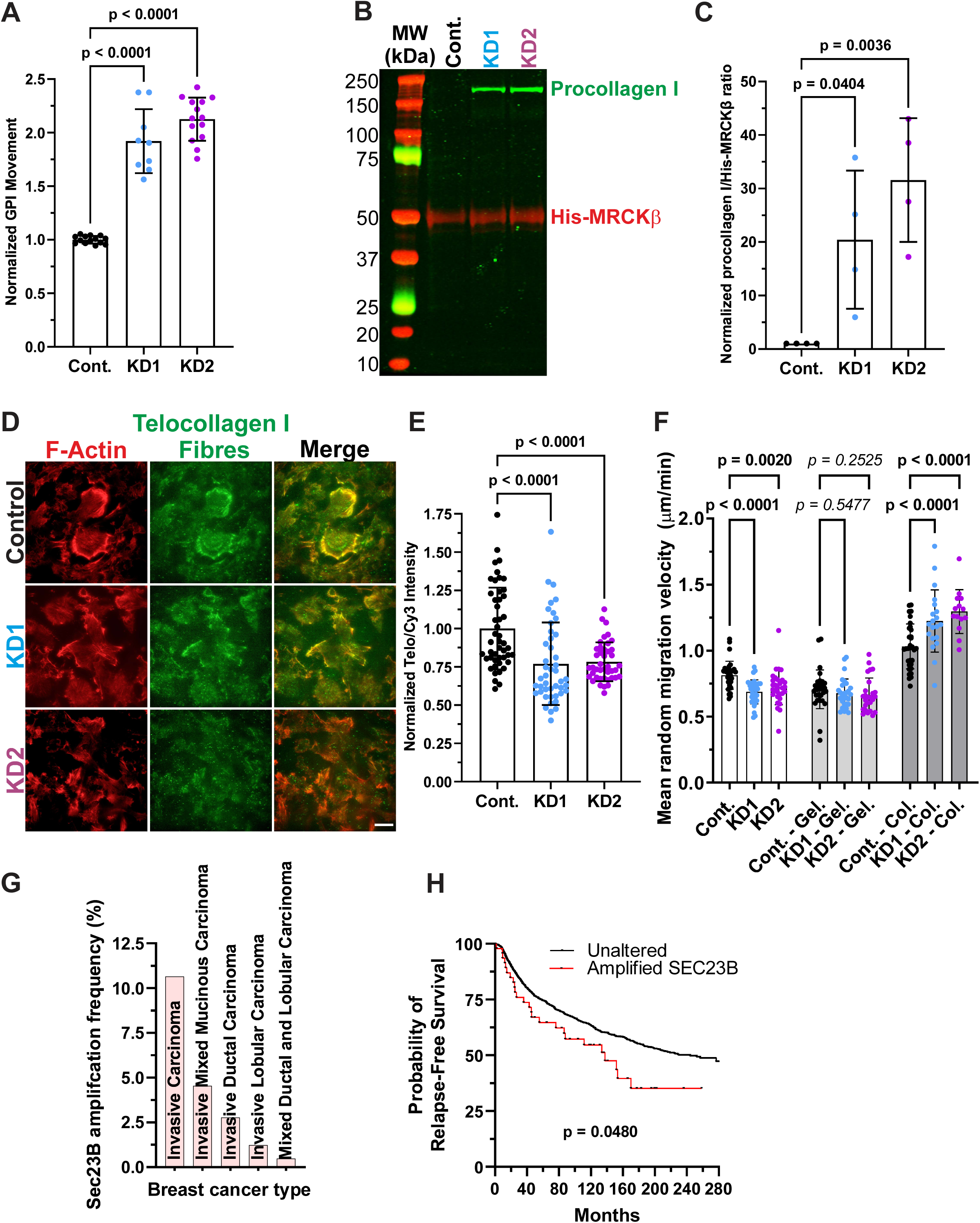
Sec23b depletion alters collagen I secretion and processing. **A.** Velocity of GPI vesicles in MDA-MB-231 cells expressing control scrambled shRNA (n = 14), or Sec23b KD1 (n = 9) or KD2 (n = 14) shRNAs. One-way ANOVA with Dunnett’s multiple comparisons test. Means ± standard deviation. **B.** Representative western blot of concentrated media from MDA-MB-231 cells expressing control scrambled shRNA or Sec23b KD1 or KD2 shRNAs blotted with anti-bodies against procollagen I or anti-His epitope tag. **C.** Normalized ratios of procollagen I to recombinant His-tagged MRCKβ control protein in conditioned media from MDA-MB-231 cells expressing control scrambled shRNA, or Sec23b KD1 or KD2 shRNAs (n = 4). One-way ANOVA with Dunnett’s multiple comparisons test. Means ± standard deviation. **D.** Immunoflourescence images of MDA-MB-231 cells, expressing control scrambled shRNA, or Sec23b KD1 or KD2 shRNAs, stained for F-Actin with Cy3-labelled fluorescent phalloidin or anti-telocollagen I antibody. **E.** Normalized ratios of telocollagen I fluorescence to Cy3-phalloidin fluorescence intensity for MDA-MB-231 cells expressing control scrambled shRNA (n = 48), or Sec23b KD1 (n = 45) or KD2 (n = 42) shRNAs. One-way ANOVA with Dunnett’s multiple comparisons test. Means ± standard deviation. **F.** Random migration velocities for MDA-MB-231 cells expressing control scrambled shRNA (n = 14), or Sec23b KD1 (n = 9) or KD2 (n = 14) shRNAs. One-way ANOVA with Dunnett’s multiple comparisons test. Means ± standard deviation. **G.** Frequency of *Sec23b* gene amplification in invasive breast cancers from 1589 patients using cBioPortal to query the METABRIC database of somatic copy number aberrations. **H.** Probability of relapse-free survival of 1962 patients divided at the median expression level into high and low expressing cohorts. Logrank test p value shown.

To determine if the decrease of deposited fibrillar collagen I was a contributory factor for the migratory defects seen in Sec23b knockdown cells in **Figures 3I-K**, we coated tissue culture surfaces with filament-forming rat tail type I collagen or gelatin, a form of collagen I fragmented by heat denaturation and acid hydrolysis that cannot form fibres. As shown in **Figures 4F and S2E**, the migration defect of MDA-MB-231 cells on untreated glass surfaces induced by Sec23b depletion was reversed when the same cells were plated on tissue culture surfaces coated with 2 μg/mL rat tail type I collagen but not with equal amount of gelatin. In fact, gelatin generally suppressed cell migration while rat tail collagen I generally promoted cell migration, supporting the notion that functional collagen polymer is necessary to support cell migration^92–94^. Nonetheless, when we compared the control and the Sec23b knocked down cells in each condition, only the cells plated on rat tail collagen I were rescued from the migratory defect associated with Sec23b depletion. Taken together, these results indicate that Sec23b depletion deregulates secretion, N-terminal processing and fibrillogenesis of collagen I in MDA-MB-231 cells, which collectively affect normal cell migration.

Examination via cBioPoral^50^ of the METABRIC database of somatic copy number aberrations (CNAs) from 2051 breast cancer patient samples^95^ revealed Sec23b amplifications in several invasive breast cancer types (**Figure 4G**), with Sec23b amplification being significantly associated with a lower probability of relapse-free survival (**Figure 4H**). Given that collagen I deposition has been associated with reduced survival, increased metastasis and resistance to therapy in breast cancer patients^96^, Sec23b may play a contributory role by enabling collagen I secretion and activation.

## Discussion

The initiation of cell migration leads to profound reorganization of the actin cytoskeleton^18^. Although extensive prior research has identified many actin-associated proteins, these studies have primarily been based on examining cells at steady-state. By linking the G- and F-actin binding peptide Lifeact with the rapid-acting miniTurbo promiscuous biotin ligase, a tool was developed that enables characterization of the dynamic changes in the actin-associated proteome. The miniTurboID-Lifeact probe was initially validated by comparing the differences in protein biotinylation in cells with intact actin cytoskeletons *versus* cells treated with the actin-polymerization inhibitor Cytochalasin D to shift the total cellular pool from F-actin to G-actin (**Figures 1B-D**). This miniTurboID-Lifeact probe was also used to identify actin-associated proteins (actin cytoskeletome) during the early stages of cell migration into scratched areas (**Figures 1G-I**). An advantage of this probe is that the expression of the probe and labelling with biotin can be minimized to pulse label the actin proximal proteins during a specific time window.

Scratch wounding confluent monolayers of MDA-MB-231 cells and then allowing 1.5 h for cells to initiate migration led to a significant ∼50% in F-actin levels (**Figure 1F**). In addition, cytoskeleton structures including filopodia, small lamellopdia and large membrane ruffles were more prominent after inducing migration by scratching. Therefore, not only do migrating cells have more actin fibres but the cytoskeleton organization is more complex than in static resting cells. The miniTurboID-Lifeact tool identified a set of proteins that were differentially biotinylated in scratched migrating cells *versus* stationary cells (**Figure 1G**). When only considering proteins with high confidence identification and changes in biotinylation that were increased or decreased by greater than 2-fold, GSEA revealed that the most prominent molecular function was “actin binding” (**Figure 1H**). Interestingly, there were parallels in the proteins that were less biotinylated in Cytochalasin D treated cells in which F-actin was disrupted and more biotinylated in scratched cells in which F-actin levels were increased, consistent with their primary function being binding to actin filaments. However, there were also proteins with known F-actin binding properties that were less biotinylated both in Cytochalasin D treated cells and in migrating cells, suggesting that their primary function may be to organize F-actin into specific structures or to induce protein-protein complexes that are less abundant in migrating cells. This protein set included including AFAP1, Coronin 2A, Coronin 2B, Drebin1, Supervillin, Synaptopodin, Caldesmon 1, Nexilin, LIMA1, Phostensin, and FGD4. For example, Caldesmon 1 is an actin-filament binding protein that inhibits myosin binding^97^, while Caldesmon 1 knockdown was reported to increase prostate cancer cell migration and invasion^98^, consistent with its reduced association with F-actin positively contributing to facilitating the initiation of cell migration. Similarly, Coronin proteins bind to F-actin and promote filament severing and de-branching by recruiting cofilin and interacting with Arp2/3 complexes, respectively^99^, thus their reduced labelling by miniTurboID-Lifeact in migrating cells likely reflects their exclusion from the cytoskeleton to enable increased F-actin stability and arborization. Thus, this miniTurboID-Lifeact probe is useful for labelling actin cytoskeletomes at a specific time and space to reveal unidentified proteins that control or associate with the actin cytoskeleton during different cellular processes.

An unexpected finding was that Sec23b, a component of COPII vesicles, was the third most highly miniTurboID-Lifeact-mediated biotinylated protein following scratch-induced migration (**Figure 1G)**. Furthermore, Sec23b was less biotinylated in Cytochalasin D treated cells and more biotinylated in scratched cells expressing either miniTurboID-Lifeact or Lifeact-miniTurboID (**Figure 2A)**, indicating that it was proximal to the F-actin cytoskeleton and independent of the arrangement of the probe functional domains. A proteomics study in which the F-actin binding protein Zyxin was coupled to the TurboID promiscuous biotin ligase identified the Sec23b partner Sec23a as being more biotinylated when actin fibres were induced by cell stretching^100^, consistent with COPII proteins being proximal to the actin cytoskeleton. An siRNA screen determined that there were functional relationships between Sec23a and Sec23b with cytoskeleton-associated proteins including Moesin and Radixin (identified as being more biotinylated in migrating cells in **Figure 1I**) that affected the transportation of cargo proteins from the ER to the cell surface^101^. Co-immunoprecipitation experiments did not detect a direct physical interaction between Sec23b and the microtubule-actin cross-linker MACF1 that had been one of the siRNA hits, raising the possibility that there may be an intermediary, such as the actin cytoskeleton, which enables the functional interaction. Sec23b knockdown did not observably affect actin structures, indicating that is unlikely to be a direct regulator of the actin cytoskeleton (**Figures 3B and 3C**). However, Sec23b knockdown did affect cell spreading, focal adhesion formation/maturation and migration (**Figures 3E-K**), consistent with it indirectly influencing these cell behaviours.

The COPII protein complex facilitates ER-to-Golgi transport thereby assists transportation of secretory proteins that are destined to their extracellular destination^33,63^. Many extracellular matrix proteins, including collagens, are packaged into COPII-coated vesicles for transport from the ER to the Golgi apparatus, where collagen I undergoes post-translational modifications (*e.g.* glycosylation) before being secreted^78,87,92,96^. For collagen I proteins, newly synthesized soluble collagen I proteins are assembled into triple-helices (procollagen I) in the ER and secreted into the extracellular matrix. The secreted soluble extracellular procollagen I must be cleaved at the N- and C-terminii by metalloproteases (*e.g.* ADAMTS, a disintegrin and a metalloproteinase with thrombospondin repeats, and BTPs, bone morphogenetic protein-1/Tolloid-like proteinases)^87^ for the collagen I to assembled into insoluble fibres. Based on the antibodies used, we deduced that the N-terminal domain of procollagen was not cleaved, thus preventing the fibrillogenesis of collagen I^92,102^. The observed increase in secreted extracellular procollagen I and concomitant decrease in fibrillar collagen I suggests an imbalance between procollagen I secretion and the secretion and/or activity of metalloproteases such as ADAMTS responsible for cleaving the procollagen I N-terminus. Alternatively, depletion of Sec23b could cause misfolding of the procollagen I protein, resulting in the N-terminal procollagen I cleavage site becoming masked from extracellular proteases. Interestingly, the observed decrease in the migration of MDA-MB-231 cells with Sec23b depletion was rescued by pre-coating the surface with rat tail collagen I but not with gelatin (**Figure 4F**). Thus, Sec23b appears to be important in balancing collagen I secretion and its post-secretion maturation for efficient migration of MDA-MB-231 cells.

Pulse-induction of the miniTurboID-Lifeact probe clearly differentially labeled proteins under varying conditions of actin cytoskeleton organization. We identified Sec23b as a factor that influenced scratch-induced and random migration in MDA-MB-231 cells by affecting the secretion and processing of collagen I. Additional proteins were identified as being associated with the cytoskeleton to greater or lesser degrees in cells undergoing early stages of cell migration, suggesting that there are more proteins with uncharacterized roles in enabling cell movement.

## Methods

### Plasmids and DNA constructs

All oligos were synthesized by Integrated DNA technologies and all restriction enzymes were purchased from New England Biolabs. mApple-GPI-RUSH was a gift from Jennifer Lippincott-Schwartz (Addgene plasmid #166904; http://n2t.net/addgene:166904; RRID:Addgene_166904)^103^, pLVXpuro-mGFP-Sec23b was a gift from David Stephens (Addgene plasmid #66598; http://n2t.net/addgene:66598; RRID:Addgene_66598)^104^, mCherry-EB3-7 was a gift from Michael Davidson (Addgene plasmid #55037; http://n2t.net/addgene:55037; RRID:Addgene_55037), pEB3-miRFP670nano was a gift from Vladislav Verkhusha (Addgene plasmid #127433; http://n2t.net/addgene:127433; RRID:Addgene_127433)^105^, pSBtet-Bla was a gift from Eric Kowarz (Addgene plasmid #60510; http://n2t.net/addgene:60510; RRID:Addgene_60510)^106^, pSBtet-Hyg was a gift from Eric Kowarz (Addgene plasmid #60508; http://n2t.net/addgene:60508; RRID:Addgene_60508)^106^, and pemiRFP670-N1 was a gift from Vladislav Verkhusha (Addgene plasmid #136556; http://n2t.net/addgene:136556; RRID:Addgene_136556)^107^. pCMV-VSV-G was a gift from Bob Weinberg (Addgene plasmid #8454; http://n2t.net/addgene:8454; RRID:Addgene_8454)^108^. psPAX2 was a gift from Didier Trono (Addgene plasmid #12260; http://n2t.net/addgene:12260; RRID:Addgene_12260). pENTR1A was purchased from Thermo Fisher Scientific. To generate N- or C-terminus miniTurboID-Lifeact, 5’-AATTatgggcgtggccgacttgatcaagaagttcgagtccatctccaaggaggag-3’ and 5’-TCGActcctccttggagatggactcgaacttcttgatcaagtcggccacgcccat-3’ were annealed then subcloned into EcoRI and XhoI sites on pENTR1A (pENTR1A-Lifeact). pENTR1A-Lifeact was subsequently Gateway LR cloned into pSTV6-N-miniTurbo-BirA (miniTurboID-Lifeact) or pSTV6-C-miniTurbo-BirA (Lifeact-miniTurboID). For control miniTurboID that lacks Lifeact amino acid segment, pENTR1A was Gateway LR cloned into pSTV6-N-miniTurbo-BirA or pSTV6-C-miniTurbo-BirA. To generate shRNA-Sec23b-KD1 (KD1), oligo pair of 5’-ccggGCCTGTTCACAAGATTGATATctcgagATATCAATCTTGTGAACAGGCttttg-3’ and 5’-aattcaaaaGCCTGTTCACAAGATTGATATctcgagATATCAATCTTGTGAACAGGC-3’ were annealed first then subcloned into AgeI and EcoRI sites on Tet-pLKO-puro. Similar oligo cloning strategies were utilized to generate shRNA-Sec23b-KD2 (KD2) and shRNA-Scrambled-Control (Control). The oligo pairs used for annealing are as follows: KD2: 5’-ccggCCACTTTGTCAGGTTGATTATctcgagATAATCAACCTGACAAAGTGGtttt-3’ and 5’-aattaaaaCCACTTTGTCAGGTTGATTATctcgagATAATCAACCTGACAAAGTGG-3’; and Scrambled control: 5’-CCGGGCGCGATAGCGCTAATAATTTCTCGAGAAATTATTAGCGCTATCGCGCTTTT-3’ and 5’-AATTCAAAAAGCGCGATAGCGCTAATAATTTCTCGAGAAATTATTAGCGCTATCGCGC-3’. pSBtet-Bla-EB3-7-670nano was generated by Gibson assembly (New England Biolab) of pEB3-miRFP670nano and pemiRFP670-N1 into pSBtet-Bla. Specifically, a primer pair of 5’-AGAGCTAAGGCCTGTCAGGCCAAGC-3’ and 5’-GCCATGGTGGTGGCCTCAGAGGCCTTT-3’ was used to generate the linearized vector (pSBtet-Bla), while a fragment containing EB3 was generated by primer pair consisting of 5’-GGCCACCACCATGGCCGTCAATG-3’ and 5’-TCCGCCATGGTGGCGACCGG-3’ and a fragment containing emiRFP670 was generated by the primer pair of 5’-CGCCACCATGGCGGAAGGC-3’ and 5’-GACAGGCCTTAGCTCTCAAGCGCGGTGA-3’. Similarly, pSBtet-Hyg-Lifeact-mCherry was constructed by Gibson assembly of pENTR1A-Lifeact and mCherry-EB3-7 into pSBtet-Hyg. Primers used to linearize the vector (pSBtet-Hyg) were 5’-ACAAGTAAGGCCTGTCAGGCCAA-3’ and 5’-ACGCCCATGGTGGCCTCAGAGG-3’.

The primers used to generate fragments containing Lifeact were 5’-GGCCACCATGGGCGTGGCCGAC-3’ and 5’-CTCACCATCTCCTCCTTGGAGATGGACTCG-3’, and the primers used to generate fragments containing mCherry were 5’-AGGAGGAGATGGTGAGCAAGGGCGAG-3’ and 5’-TGACAGGCCTTACTTGTACAGCTCGTCCATGCC-3’.

### Cell Culture, cell lines, and transfection

Human embryonic kidney cells (HEK293Tv), metastatic breast cancer cells (MDA-MB-231) and their derivative stable cell lines were grown in DMEM (Sigma-Aldrich) supplemented with 10% heat-inactivated FBS (Gibco) at 37°C and 5% CO_2_ in a humid environment. For generating stable cell lines, lentiviral particles were packaged in HEK 293Tv cells by co-transfecting miniTurboID-Lifeact, Lifeact-miniTurboID, pSTV6-N-miniTurbo-BirA, pSTV6-C-miniTurbo-BirA, shRNA-Sec23b-KD1, shRNA-Sec23b-KD2, or shRNA-Scramble-Control or with pLVXpuro-mGFP-Sec23b with psPAX2 and pVSVG using PolyJet in Vitro DNA transfection reagent according to the manufacturer’s instructions (SignaGen). Media were replaced with fresh media 24 h post-transfection and were collected 48 h post-transfection. The collected media were passed through a 0.45 μm filter and then added to 60% confluent MDA-MB-231 cells. Media containing viral particles were removed 24 h post infection and cells were washed twice with fresh media. Stably transfected cells were selected 72 hours post-infection with 2 μM puromycin (Sigma-Aldrich). For generating cell line stably expressing fluorophore tagged Sec23b, Lifeact and EB3, MDA-MB-231 cells stably expressing pLVXpuro-mGFP-Sec23b were sequentially transfected with pSBtet-Hyg-Lifeact-mCherry then with pSBtet-Bla-EB3-7-670nano using an Amaxa 4D-nucleofector with SE cell line kit (Lonza). Three days after the first transfection, cells were selected with 50 μM hygromycin for two weeks and selected with 2 μM Blasticidin three days after the second transfection. For enriching cells expressing low levels of all tagged Sec23b, Lifeact and EB3, cells were sorted using fluorescence-activated cell sorting (BD Biosciences).

### BioID sample preparation and mass spectrometry

To perform BioID^109^, stable cells expressing miniTurboID-Lifeact, Lifeact-miniTurboID or the control miniTurboID were induced with 1 μg/ml of doxycycline (Sigma-Aldrich) for 18 h at 37°C and 5% CO_2_ followed by 30 min incubation with 50 mM Biotin (BioShop) before collecting the cells. For cytochalasin D condition, cells were treated with 5 μM cytochalasin D (BioShop) for 1 h before labelling with biotin. To label proteins at the early stages of scratch-induced invasion/migration, ∼90% confluent cells were scratched 1 mm apart vertically and horizontally and then washed twice with PBS. These cells were allowed to migrate/invade into the scratched area for 1 h at 37°C and 5% CO_2_ before labelling with 1 mM of biotin for 30 min. Cells were collected by centrifuging at 2000 rpm for 3 minutes. The pelleted cells were washed twice with PBS and then snap frozen. The frozen cell pellets were processed for mass spectrometry as described previously^110^. Briefly, the collected cells were pelleted (2000 rpm, 3 min), washed twice with PBS and saved by snap freezing. The cell pellet was resuspended in 10 mL of lysis buffer (50 mM Tris-HCl pH 7.5, 150 mM NaCl, 1 mM EDTA, 1 mM EGTA, 1% Triton X-100, 0.1% SDS) supplemented with 1:500 protease inhibitor cocktail (Sigma-Aldrich) and 1:1,000 benzonase nuclease (Novagen) and incubated on an end-over-end rotator at 4°C for 1 h, briefly sonicated to disrupt visible aggregates, then centrifuged at 45,000 X g for 30 min at 4°C. The supernatant was transferred to a fresh 15 mL conical tube. 30 mL of packed, pre-equilibrated Streptavidin-Sepharose beads (GE) were added and the mixture was incubated for 3 hours at 4°C with end-over-end rotation. Beads were pelleted by centrifugation at 2000 rpm for 2 min and transferred with 1 mL of lysis buffer to a fresh Eppendorf tube. Beads were washed once with 1 mL lysis buffer and twice with 1 mL of 50 mM ammonium bicarbonate (pH 8.3). Beads were transferred in ammonium bicarbonate to a fresh centrifuge tube and washed twice with 1 mL ammonium bicarbonate buffer. Tryptic digestion was performed by incubating the beads with 1 mg MS-grade TPCK trypsin (Promega) dissolved in 200 mL of 50 mM ammonium bicarbonate (pH 8.3) overnight at 37° C. The following morning, 0.5 mg MS-grade TPCK trypsin was added, and beads were incubated for 2 additional hours at 37°C. Beads were pelleted by centrifugation at 2,000 X g for 2 min, and the supernatant was transferred to a fresh Eppendorf tube. Beads were washed twice with 150 mL of 50 mM ammonium bicarbonate, and these washes were pooled with the first eluate. The sample was lyophilized and resuspended in buffer A (0.1% formic acid). 1/5th of the sample was analyzed per MS run.

Mass spectrometry of BioID samples was conducted on high-performance liquid chromatography using a 2 cm pre-column (Acclaim PepMap 50 mm x 100 µm inner diameter (ID)), and 50 cm analytical column (Acclaim PepMap, 500 mm x 75 µm diameter; C18; 2 µm; 100 Å, Thermo Fisher Scientific), running a 120 min reversed-phase buffer gradient at 225 nL/min on a Proxeon EASY-nLC 1000 pump in-line with a Thermo Q-Exactive HF quadrupole-Orbitrap mass spectrometer. A parent ion scan was performed using a resolving power of 60,000, and then up to the twenty most intense peaks were selected for MS/MS (minimum ion count of 1,000 for activation) using higher energy collision induced dissociation (HCD) fragmentation. Dynamic exclusion was activated such that MS/MS of the same m/z (within a range of 10 ppm; exclusion list size = 500) detected twice within 5 s were excluded from analysis for 15 s. For protein identification, Thermo.RAW files were converted to the.mzXML format using Proteowizard^111^, then searched using X!Tandem^112^ and COMET^113^ against the Human RefSeq Version 45 database (containing 36,113 entries). Data were analyzed using the trans-proteomic pipeline (TPP)^114,115^ via the Pro-Hits software suite (v3.3)^116^. Search parameters specified a parent ion mass tolerance of 10 ppm, and an MS/MS fragment ion tolerance of 0.4 Da, with up to 2 missed cleavages allowed for trypsin. Variable modifications of +16@M and W, +32@M and W, +42@N-terminus, and +1@N and Q were allowed. Proteins identified with an iProphet cut-off of 0.9 (corresponding to %1% FDR) and at least two unique peptides were analyzed with SAINT Express v.3.6. Twenty control runs (from cells expressing the miniTurboID* epitope tag) were collapsed to the two highest spectral counts for each prey and compared to the two biological replicates (each with two technical replicates) of miniTurboID-Lifeact. High confidence interactors were defined as those with Bayesian false discovery rate (BFDR) % 0.05.

### Immunofluorescence, live cell imaging, and analysis

Stable cells that were pre-treated with doxycycline were plated on 1.5-thickness glass coverslips (EMS) and grown for at least 24 h. Cells were fixed for 15 min at room temperature with 4% paraformaldehyde in PBS supplemented with 5% sucrose (BioShop) followed by 0.1% TX-100 (Sigma-Aldrich) solubilization for 3-5 min at room temperature. Fixed cells were blocked with 30% normal donkey serum (Jackson Immunoresearch) supplemented with 5% BSA (Sigma Aldrich) for 1 h at room temperature, then incubated with primary antibody for 12-18 h at 4°C. After washes in PBS, cells were incubated with secondary antibodies, phalloidin conjugated with Rhodamine or Alexa Fluor 488 (Life Technology), and/or streptavidin conjugated with Alexa Fluor 647 (Life Technology). The following antibodies were used: rabbit anti-Sec23b (Abcam; 1:1000), rabbit anti-telocollagen (Antibodies Inc; 1:200), mouse anti-FLAG (Sigma Aldrich; 1:200), mouse anti-paxillin (BD Biosciences; 1:400); and secondary antibodies conjugated with Alexa Fluor488, 568, or 647 (Jackson Immunoresearch). Fixed and immunostained cells on coverslips were visualized using a 63x/1.49 NA TIRF objective on Quorum Diskovery microscope system equipped with a Leica DMi8 microscopy connected to a Zyla 4.2Plus sCMOS camera and controlled by Quorum WaveFX powered by MetaMorph software (Quorum Technologies). Paxillin positive adhesion sizes were measured as described in^117^ after deconvolving images with Huygens Professional version 23.04 (Scientific Volume Imaging, The Netherlands). Elongation factor was measured by dividing width by length (MetaMorph, Molecular Devices).

Live-cell imaging was performed with a Quorum Diskovery Spinning Disc Confocal microscope system equipped with environmental chambers set to 37°C and 5% CO_2_ and with a Leica DMi8 microscopy connected to an iXon Ultra 897 EMCCD BV camera and controlled by Quorum WaveFX powered by MetaMorph software (Quorum Technologies). FACS-sorted cells were induced first with 50 nM doxycycline for 12-18 hours. When necessary, scratches were generated, followed by 3 washes with pre-warmed complete media, then incubated in the environmental chamber for one hour, and imaging was completed within 30 minutes. For measuring the movements of GPI-marked vesicles, RUSH (retention using selective hooks) assays were performed^118,119^. Briefly, mApple-GPI-RUSH was expressed in Sec23b depleted or control cells and the release of GPI-marked cargos was synchronized by incubating with 80 μM biotin. Imaging was recorded within 45 minutes. Sec23b vesicle movements were analyzed by Particle Tracker, a part of the MosaicSuite^120,121^.

### Wound healing and migration studies

For random migration studies, appropriate stable cells were induced with 2 μM doxycycline for 2-3 days, then replaced with fresh complete media (media supplemented with 10% FBS and 2 μM doxycycline) every 24 hr. On the day before experiments, cells were plated on Essen Bioscience ImageLock 96-well plates (Sartorius) at 1500 cells per well and incubated 12-18 h at 37°C 5% CO_2_ before replacing with fresh complete media and imaging in an IncuCyte S3 chamber (Sartorius) every 10 minutes for 12-18 h. For the ECM rescue studies, Essen Bioscience ImageLock 96-well plates were pre-coated with the indicated ECM for 1 h at 37°C 5% CO_2_, then washed three times with PBS before plating cells. Cells were incubated in the IncuCyte S3 chamber for 2 h and imaged as described above. Images were exported as MetaMorph-compatible stacks, and at least 6 h of movement were analyzed using the MetaMorph (Molecular Devices) automatic tracking app. More than 123 cells were analyzed from 6 independent wells per condition.

Similarly, for scratch-induce migration studies, cells were induced with 2 μM doxycycline for 2-3 days, replacing with fresh complete media (media supplemented with 2 μM doxycycline) for every 24 h. On the following day, cells were plated on Essen Bioscience ImageLock 96-well plates at 4 x 10^4^ cells per well and incubated 12-18 h at 37°C 5% CO_2_ to ensure a confluent monolayers cells at the end of the incubation. A uniform scratch was generated using the IncuCyte Woundmaker Tool (Sartorius) as instructed by the manufacturer. Cells were washed three times with complete media and then imaged in an IncuCyte S3 chamber every 15-20 min for 48 h. Scratched width was measured using IncuCyte S3 software.

### Secretion assays

Protein samples were either extracted from whole cells by lysing in 2X complete LDS sample buffer (Thermo Scientific) or concentrated from media with StrataClean resin (Agilent Technologies) as described previously^122^. Briefly, cells were plated on 6-well dishes and induced with 2 μg/mL doxycycline for 3 days, replacing with fresh complete media (media supplemented with 10% FBS and 2 μM doxycycline) for every 24 h. On the fourth day, cells were washed 5X with DMEM supplemented only with 2 μg/ml doxycycline (Sigma-Aldrich) then incubated with 1 mL of doxycycline supplemented DMEM for 48 hours. At the end of incubation, media was collected and spun at 5000 rpm for 10 min at 4°C. Cleared media were incubated with pre-treated StrataClean resin and the mixture incubated for overnight at 4°C with end-over-end rotation.

### Western blotting

Samples were subjected to SDS-PAGE followed by a transfer onto nitrocellulose membranes (BioRad). Blots were blocked with 1% skim milk for 1 h at room temperature then incubated with primary antibodies for 1 h at room temperature or overnight at 4°C. After washes in PBS, blots were subjected to secondary antibodies for 1 h at room temperature then detected using an Odyssey infrared imaging system (LI-COR). The following antibodies were used: rabbit anti-Sec23b (Abcam; 1:1000); mouse anti-FLAG (Sigma-Aldrich; 1:1000); mouse anti-vimentin (Sigma-Aldrich; 1:2000); anti-GAPDH (DHSB; 1:2500); anti-procollagen clone SP1.D8 (DHSB; 1:200); rabbit anti-His (Cell Signaling Tech; 1:1000); secondary antibodies conjugated to IRDye 680 or 800 (LI-COR).

### Cell spreading and analysis

Cells depleted of Sec23b or corresponding controls were plated on Essen Bioscience ImageLock 96-well plates at 2000 cells per well and immediately imaged every 5 minutes for 6 hours at 20X magnification. Images were exported as MetaMorph compatible stacks and areas of cells were measured with MetaMorph (Molecular Devices).

### Statistics

All statistics were performed with GraphPad Prism 10.6.1 and reported in figure legends.

## Funding

This study was supported by the Philip S. Orsino Chair in Leukemia Research, a joint Hospital-University Named Chair between the University of Toronto, The Princess Margaret Cancer Centre Director, and the Princess Margaret Cancer Foundation (B.R.), the Canada Research Chairs program (950-231665, CRC-2024-00113; M.F.O.) and Natural Sciences and Engineering Council of Canada (RGPIN-2020-05388; M.F.O).

### Author contributions

E.E.J. and M.F.O. conceptualized, designed the research work and established the methodology. E.E.J. and A.A. investigated. A.A. assisted in the creation of stable cell lines, and generated the samples for mass spectrometry. J.S.G. and B.R. collected and curated the mass spectrometry data. E.E.J. and M.F.O. analyzed the data and performed statistical analyses. M.F.O. administered, supervised and acquired funding for the project. E.E.J. wrote the first draft. M.F.O led the preparation of the final version with the involvement of all authors.

## Supporting information

Supplemental figures and titles

