## Supplemental figures and titles for "Sec23b regulates cell migration by orchestrating collagen I secretion and processing"

**Supplemental Information:** **Page**

**Figure S1. No effect of miniTurboID-Lifeact on cell migration.** 2

**Figure S2. Cell migration parameters.** 4

**Supplemental Table S1. Proteins labelled by miniTurboID-Lifeact affected by Cytochalasin D.** 6

**Supplemental Table S2. GSEA molecular function gene sets with increased Cytochalasin D induced biotinylation by miniTurboID-Lifeact.** 6

**Supplemental Table S3. GSEA molecular function gene sets with deceased Cytochalasin D induced biotinylation by miniTurboID-Lifeact.** 6

**Supplemental Table S4. Proteins labelled by miniTurboID-Lifeact affected by scratch wounding.** 6

**Supplemental Table S5. GSEA molecular function gene sets with increased scratch wound induced biotinylation by miniTurboID-Lifeact.** 6

**Supplemental Table S6. GSEA molecular function gene sets with decreased scratch wound induced biotinylation by miniTurboID-Lifeact.** 6

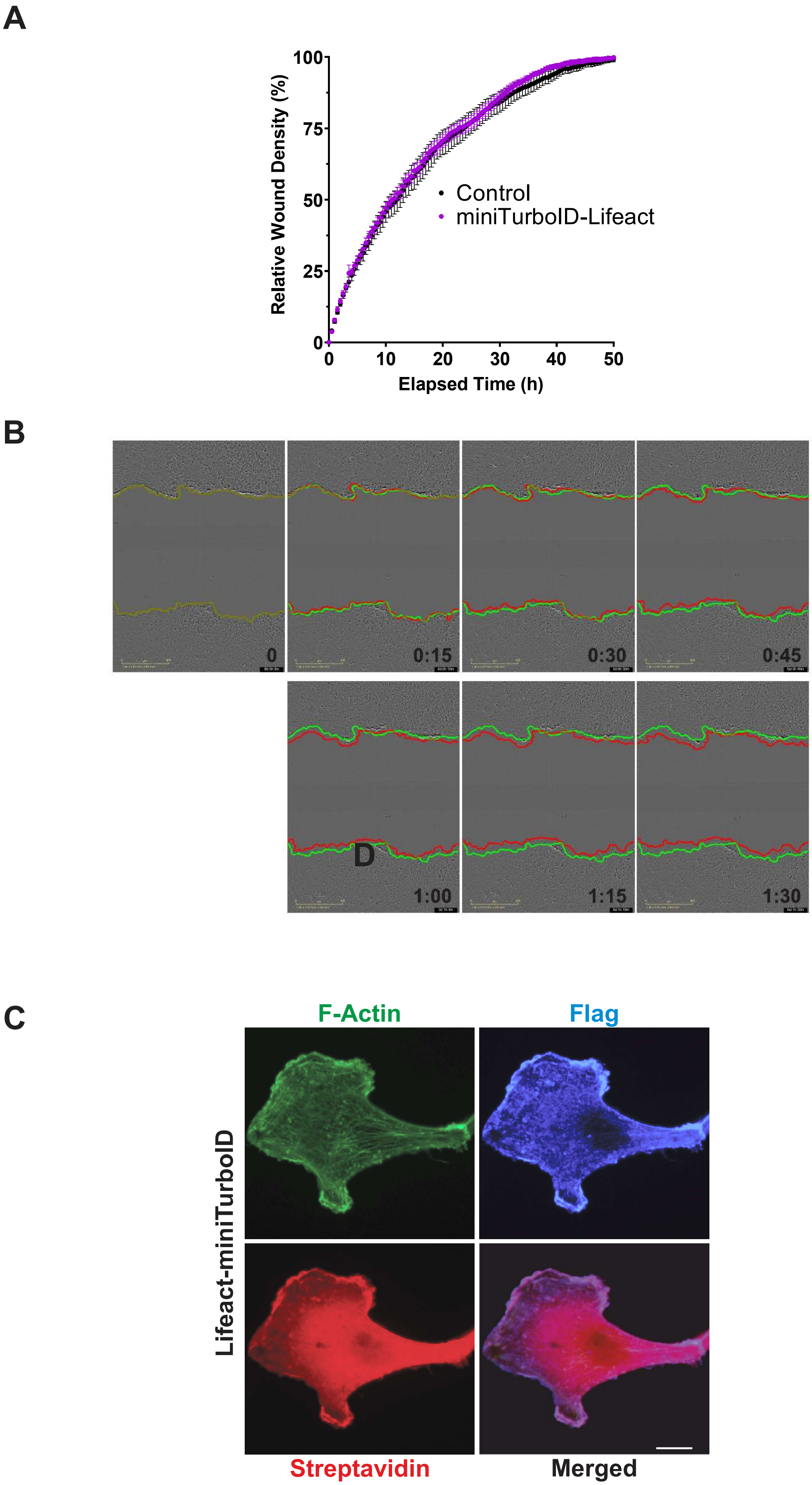
A
100
75
50
25
0
0
10
20
30
40
50
Relative Wound Density (%)
Elapsed Time (h)
Control
miniTurboID-Lifeact
B
0
0:15
0:30
0:45
D
1:00
1:15
1:30
C
F-Actin
Flag
Lifeact-miniTurboID
Streptavidin
Merged

Figure S1

**Figure S1. No effect of miniTurboID-Lifeact on cell migration.**

**A.** Scratch induced cell migration was not different between MDA-MB-231 cells in which expression of miniTurboID-Lifeact was uninduced (Control) or induced. **B.** Time course of scratch wound induced cell migration shows that cells had begun to move into the cell-free scratched area in 1.5 h (red lines) relative to their starting positions (green lines). Scale = 300 $\mu$m. **C.** Example MDA-MB-231 cells expressing Lifeact-miniTurboID stained for F-actin structures with phalloidin (green), FLAG epitope tag (blue), and protein biotinylation with streptavidin (red).

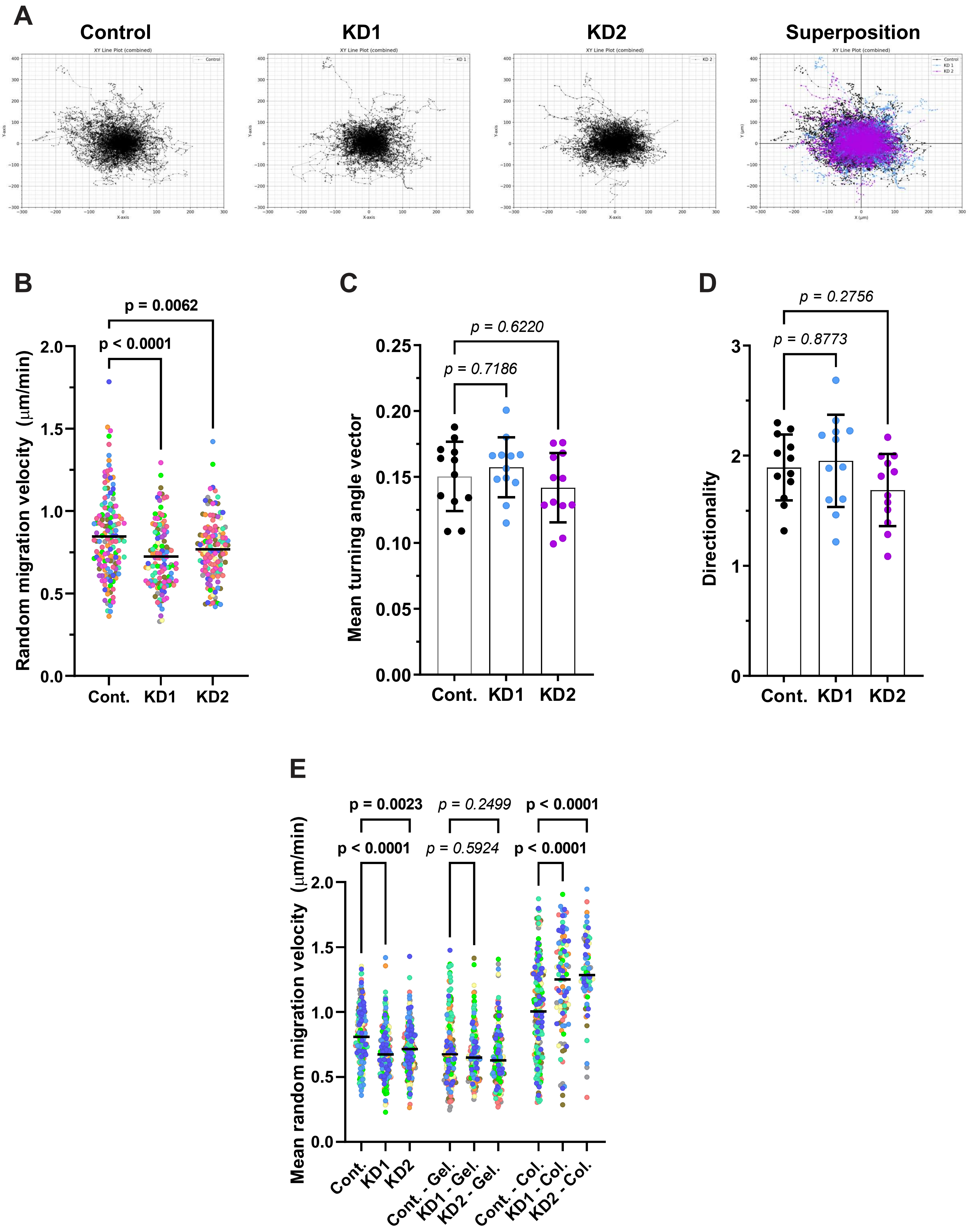
A
Control
KD1
KD2
Superposition
XY Line Plot (combined)
B
p = 0.0062
p < 0.0001
Random migration velocity (µm/min)
Cont.
KD1
KD2
C
p = 0.6220
p = 0.7186
Mean turning angle vector
Cont.
KD1
KD2
D
p = 0.2756
p = 0.8773
Directionality
Cont.
KD1
KD2
E
p = 0.0023
p = 0.2499
p < 0.0001
p < 0.0001
p = 0.5924
p < 0.0001
Mean random migration velocity (µm/min)
Cont.
KD1
KD2
Cont. - Gel.
KD1 - Gel.
KD2 - Gel.
Cont. - Col.
KD1 - Col.
KD2 - Col.

Figure S2

**Figure S2. Cell migration parameters.**

**A.** Single cell migration X-Y tracking "spider" plots for control, KD1 and KD2 cells. Superposition figure shows tracks for control (black), KD1 (blue) and KD2 (purple) cells. **B.** Single cell random migration velocity data pooled from 12 independent replicate experiments are colour coded to represent each replicate for control (n = 146), KD1 (n = 123) and KD2 (n = 130) conditions. One-way ANOVA with Dunnett's multiple comparisons test. Means indicated with a solid line. **C.** Mean random migration turning angle vector data from 12 independent replicate experiments. One-way ANOVA with Dunnett's multiple comparisons test. Means indicated with a solid line. **D.** Mean random migration directionality data from 12 independent replicate experiments. One-way ANOVA with Dunnett's multiple comparisons test. Means indicated with a solid line. **E.** Single cell random migration velocity data pooled from 9 independent replicate experiments are colour coded to represent each replicate for control (uncoated n = 199, gelatin n = 180, collagen n = 213), KD1 (uncoated n = 198, gelatin n = 160, collagen n = 96) and KD2 (uncoated n = 192, gelatin n = 154, collagen n = 73) conditions. One-way ANOVA with Dunnett's multiple comparisons test. Means indicated with a solid line.

**Supplemental Table S1. Proteins labelled by miniTurboID-Lifeact affected by Cytochalasin D.**

Related to Figure 1B. Identified proteins from control and Cytochalasin D treated conditions that had Bayesian false discovery rates < 0.05. Values indicate the change in untreated/Cytochalasin D spectra ratio ($log_2$).

**Supplemental Table S2. GSEA molecular function gene sets with increased Cytochalasin D induced biotinylation by miniTurboID-Lifeact.**

Related to Figure 1C.

**Supplemental Table S3. GSEA molecular function gene sets with deceased Cytochalasin D induced biotinylation by miniTurboID-Lifeact.**

Related to Figure 1C.

**Supplemental Table S4. Proteins labelled by miniTurboID-Lifeact affected by scratch wounding.**

Related to Figure 1G. Identified proteins from control resting and scratch wound induced migrating conditions that had Bayesian false discovery rates < 0.05. Values indicate the normalized resting/scratched spectra ratio ($log_2$).

**Supplemental Table S5. GSEA molecular function gene sets with increased scratch wound induced biotinylation by miniTurboID-Lifeact.**

Related to Figure 1H.

**Supplemental Table S6. GSEA molecular function gene sets with decreased scratch wound induced biotinylation by miniTurboID-Lifeact.**

Related to Figure 1H.
